# RNA G-quadruplexes and calcium ions synergistically induce Tau phase transition *in vitro*

**DOI:** 10.1101/2024.03.01.582861

**Authors:** Yasushi Yabuki, Kazuya Matsuo, Ginji Komiya, Kenta Kudo, Karin Hori, Susumu Ikenoshita, Yasushi Kawata, Tomohiro Mizobata, Norifumi Shioda

**Affiliations:** Department of Genomic Neurology, Institute of Molecular Embryology and Genetics (IMEG), Kumamoto University, Kumamoto, Japan; Graduate School of Pharmaceutical Sciences, Kumamoto University, Kumamoto, Japan; Department of Neurology, Graduate School of Medical Sciences, Kumamoto University, Kumamoto, Japan; Department of Chemistry and Biotechnology, Graduate School of Engineering Tottori University, Tottori, Japan

**Author notes:** Correspondence should be addressed to Norifumi Shioda and Yasushi Yabuki, Norifumi Shioda, Department of Genomic Neurology, Institute of Molecular Embryology and Genetics, Kumamoto University, 2-2-1 Honjo, Chuo-ku, Kumamoto 860-0811, Japan., Yasushi Yabuki, Department of Genomic Neurology, Institute of Molecular Embryology and Genetics, Kumamoto University, 2-2-1 Honjo, Chuo-ku, Kumamoto 860-0811, Japan.

**Keywords:** RNA G□quadruplex, calcium ions, Tau, liquid-liquid phase separation, liquid–solid phase transition

## Abstract

Tau aggregation is a defining feature of neurodegenerative tauopathies, including Alzheimer’s disease, corticobasal degeneration, and frontotemporal dementia. This aggregation involves the liquid–liquid phase separation (LLPS) of Tau, followed by its sol–gel phase transition, representing a crucial step in aggregate formation both *in vitro* and *in vivo*. However, the precise cofactors influencing Tau phase transition and aggregation under physiological conditions (e.g., ion concentration and temperature) remain unclear. In this study, we unveil that nucleic acid secondary structures, specifically RNA G-quadruplexes (rG4s), and calcium ions (Ca^2+^) synergistically facilitated the sol–gel phase transition of human Tau under mimic intracellular ion conditions (140 mM KCl, 15 mM NaCl, and 10 mM MgCl_2_) at 37□ *in vitro*. In the presence of molecular crowding reagents, Tau formed stable liquid droplets through LLPS, maintaining fluidity for 24 h under physiological conditions. Notably, cell-derived RNA promoted Tau sol–gel phase transition, with G4-forming RNA emerging as a crucial factor. Surprisingly, polyanion heparin did not elicit a similar response, indicating a distinct mechanism not rooted in electrostatic interactions. Further exploration underscored the significance of Ca^2+^, which accumulate intracellularly during neurodegeneration, as additional cofactors in promoting Tau phase transition after 24 h. Importantly, our findings demonstrate that rG4s and Ca^2+^ synergistically enhance Tau phase transition within 1 h when introduced to Tau droplets. In conclusion, our study illuminates the pivotal roles of rG4s and Ca^2+^ in promoting Tau aggregation under physiological conditions *in vitro*, offering insights into potential triggers for tauopathy.

## Introduction

The intrinsically disordered protein Tau is a neuronal microtubule-binding protein encoded by the *MAPT* gene [1]. Tau mediates tubulin polymerization and microtubule stability, helping stabilize axons in the central nervous system [2, 3]. In pathological conditions, Tau dissociates from microtubules and forms aggregates as neurofibrillary tangles (NFTs), which may cause tauopathies such as Alzheimer’s disease (AD), corticobasal degeneration, and frontotemporal dementia [4, 5]. Post-translational modifications of Tau, such as phosphorylation and acetylation, bring about its detachment from microtubules via conformational changes, inducing its aggregation [3, 6]. However, the mechanisms underlying the aggregation of dissociated Tau proteins remain elusive.

Liquid-liquid phase separation (LLPS) is a common phenomenon resulting from weak multivalent interactions, leading to the formation of droplets, hydrogels, and aggregates [7-9]. Tau has a strong ability to undergo LLPS, which enhances physiological polymerization [10, 11]. Tau protein LLPS can also initiate its liquid–solid phase transition, resulting in its aggregation [11-15]. Polyanionic conditions, such as the presence of heparin or RNAs and hyperphosphorylation, promote Tau liquid–solid phase transition by enhancing LLPS [12, 14, 16, 17], suggesting that electrostatic interactions are critical for Tau condensates. However, *in vitro* experiments on Tau LLPS have been designed under non-physiological conditions, such as low salt concentrations, room temperature, and high-frequency agitation, which do not accurately represent the intracellular environment; therefore, the triggering factors contributing to the Tau phase transition remain unclear.

Here, we show that a nucleic acid secondary structure, RNA G-quadruplexes (rG4s) [18], and calcium ions (Ca^2+^) synergistically promote Tau phase transition under physiological conditions *in vitro*. Purified human Tau proteins formed liquid droplets *via* LLPS and remained in the fluid for 24 h. We found that rG4-forming RNA and Ca^2+^ promoted Tau phase transition within 24 h. Furthermore, when both rG4-forming RNA and Ca^2+^ were added to the Tau liquid droplets, a synergistic effect caused earlier phase transition and aggregation. These findings suggest that both rG4s and Ca^2+^ are important cofactors that promote the Tau phase transition.

## Results

### Cell-derived RNAs undergo Tau liquid–solid phase transition

First, we investigated whether purified human Tau undergoes LLPS *in vitro* under mimic intracellular ion conditions (140 mM KCl, 15 mM NaCl, and 10 mM MgCl_2_) at 37 □, with the molecular crowding agent polyethylene glycol (PEG). To visualize Tau condensates using fluorescence, non-labeled and fluorescein-labeled human Tau were mixed at a ratio of 1:1. Under mimic intracellular ion conditions, purified human Tau (0.5 mg/mL; 10.9 μM) underwent LLPS in the presence of > 5% PEG (**Fig. 1A**). However, in the presence of 10% PEG, the threshold Tau concentration for LLPS (**Fig. 1A**) was lowered, therefore, its LLPS is dependent on molecular crowding conditions. To characterize the dynamic nature of the droplets, we performed a fluorescence recovery after photobleaching (FRAP) assay. No significant differences were observed in Tau droplets with 10% PEG incubated at 37 □ from 1L24 h (**Fig. 1B**). We confirmed that Tau droplets induced sol-gel phase transition within 24 h under non-physiological conditions (e.g., low salt concentrations, room temperature [25_±_ 2 □]), and agitation (**Fig. S1**) [11, 14, 19]. These results suggest that molecular crowding alone is not sufficient to induce Tau phase transition under physiological conditions (**Fig. 1B**).

**Fig. 1.**
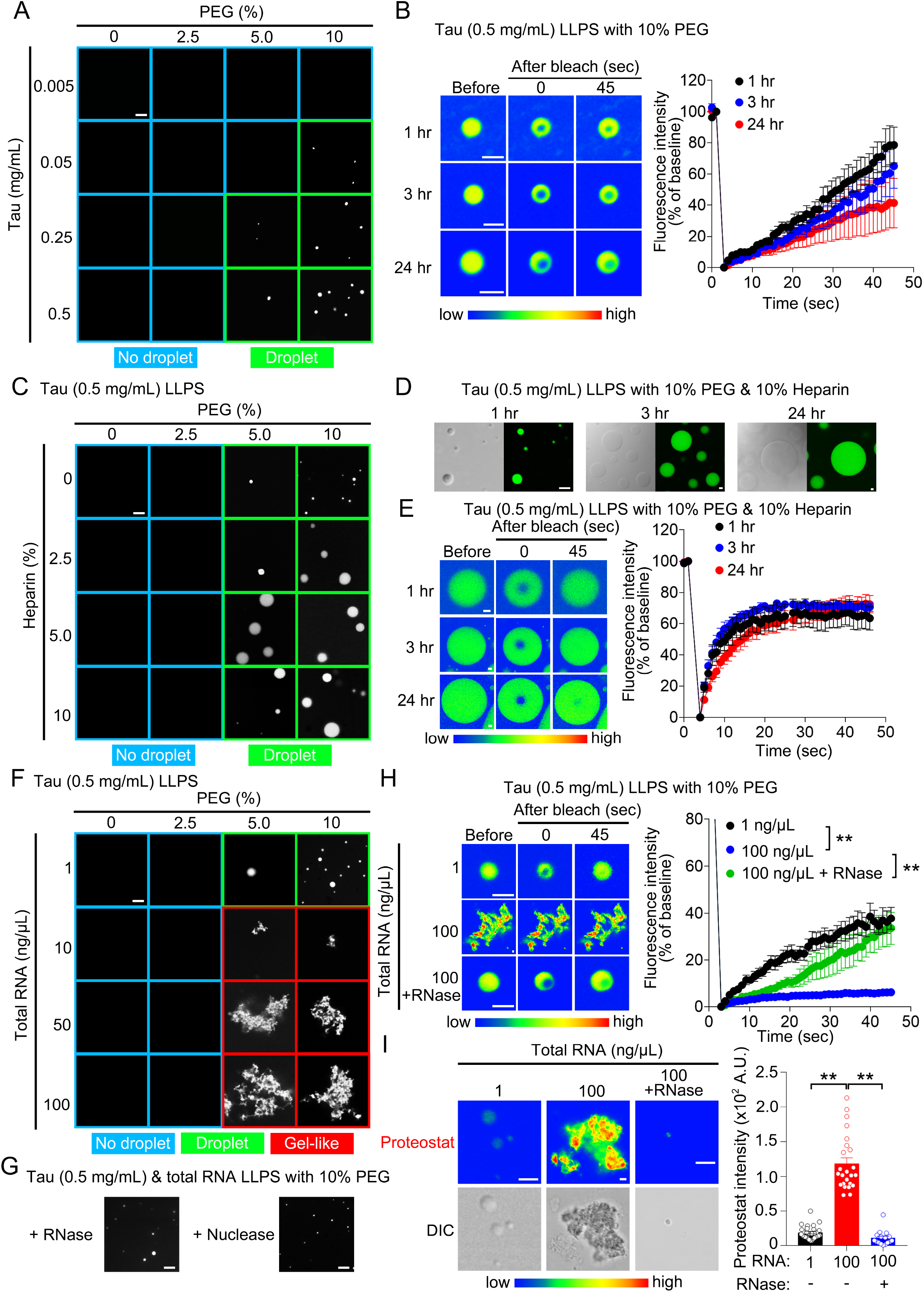
RNA initiates Tau liquid–solid phase transition. (A) Representative images of *in vitro* Tau phase separation dependent on PEG and Tau protein concentration, when incubated at 37L for 1 h. Scale bar, 5 μm. (B) FRAP assays of Tau LLPS with 10% PEG at 1, 3, and 24 h after incubation. Scale bar, 2 μm. n = 6 (1 h), n = 9 (3 h), and n = 9 (24 h). (C) Representative images of *in vitro* Tau (0.5 mg/mL) phase separation dependent on PEG and heparin concentration when incubated at 37L for 1 h. Scale bar, 5 μm. (D) Representative images of *in vitro* Tau (0.5 mg/mL) phase separation with 10% PEG and 10% heparin at 1, 3, and 24 h after incubation. Scale bar, 5 μm. (E) FRAP assays of Tau LLPS with 10% PEG and 10% heparin at 1, 3, and 24 h after incubation. Scale bar, 5 μm. n = 6 (1 h), n = 7 (3 h), and n = 6 (24 h). (F) Representative images of *in vitro* Tau (0.5 mg/mL) phase separation dependent on PEG and total RNA concentration at 1 h after incubation. Scale bar, 5 μm. (G) Representative images of *in vitro* Tau (0.5 mg/mL) phase separation with 10% PEG and 100 ng/μL total RNA in the presence of RNase A (right) and Nuclease (left) incubated at 37L for 1 h. Scale bar, 5 μm. (H) FRAP assays of Tau LLPS with 10% PEG and total RNA (1 or 100 ng/μL) with or without RNase A at 1 h after incubation. Scale bar, 2 μm. n = 5 (1 ng/μL), n = 7 (100 ng/μL), and n = 6 (100 ng/μL + RNase A). (I) Analysis of Proteostat intensity within Tau LLPS with 10% PEG and total RNA (1 or 100 ng/μL) with or without RNase A at 1 h after incubation. Scale bar, 2 μm. n = 30 (1 ng/μL), n = 24 (100 ng/μL), and n = 16 (100 ng/μL + RNase A). Data are presented as the mean ± standard error of the mean. \*\**P* < 0.01 by two-way (B, E, and H) and one-way (I) analysis of variance with Bonferroni’s multiple comparisons test.

Under the same physiological conditions, we investigated the effect of the polyanionic reagents (i.e., heparin and RNAs) on Tau LLPS. Tau droplets appeared to increase in number with increasing heparin concentrations (**Fig. 1C**). Heparin (10%) seemed to result in the time-dependent enlargement of Tau droplets without affecting fluorescence recovery (**Fig. 1D and 1E**). Tau droplets in 10% heparin showed rapid fluorescence recovery and maintained molecular diffusion inside condensates within 24 h, suggesting that polyanionic heparin may help buffer Tau inside droplets under physiological conditions. Conversely, total RNAs derived from neuro2a cells elicited Tau phase transition from a droplet to an aggregate state when incubated at 37 □ for 1 h (**Fig. 1F**). We confirmed that pretreatment with RNase A, or a nuclease, inhibited Tau phase transition (**Fig. 1G**). In addition, the application of RNA to Tau significantly reduced the fluorescence recovery of Tau condensates in FRAP and increased proteostatic intensity, which was prevented by treatment with RNase A (**Fig. 1H and 1I**).

### rG4 promotes Tau liquid–solid phase transition

Since G4-forming RNAs (rG4) have an important role in the phase transition of prion-like proteins *in vitro* and *in vivo*, including α-synuclein [20, 21], we investigated whether RNA structures are involved in Tau phase transition using various RNA oligonucleotides: G4tr, a typical rG4, telomeric repeat-containing RNA (TERRA) (UUAGGG)_4_ repeats; G4mt, a TERRA mutant that is unable to form rG4 (UUACCG)_4_ repeats, a mismatch-hairpin formed (CAG)_8_ repeats, and polyA; single-stranded (AAA)_8_ repeats [21]. Similar to other prion-like proteins, G4tr triggered a Tau phase transition, leading to an aggregate-like state over time, whereas the other RNA structures showed an increase in liquid droplet size and remained in a liquid state similar to heparin (**Fig. 2A**). In the FRAP analysis, the fluorescence recovery of G4tr-treated Tau was markedly reduced in a time-dependent manner (**Fig. 2B**). After 24 h, the fluorescence recovery rate of Tau treated with G4tr was significantly lower than that of the untreated control and G4mt-treated Tau (**Fig. 2C**). Moreover, the intensity of proteostat signals was significantly higher in G4tr-treated Tau than in Tau alone and non-G4 forming RNAs-treated Tau (**Fig. 2D**). We also confirmed the colocalization of Cyanine5-labeled G4tr with Tau droplets after 1 h and Tau aggregates after 24 h **(Fig. 2E**).

**Fig. 2.**
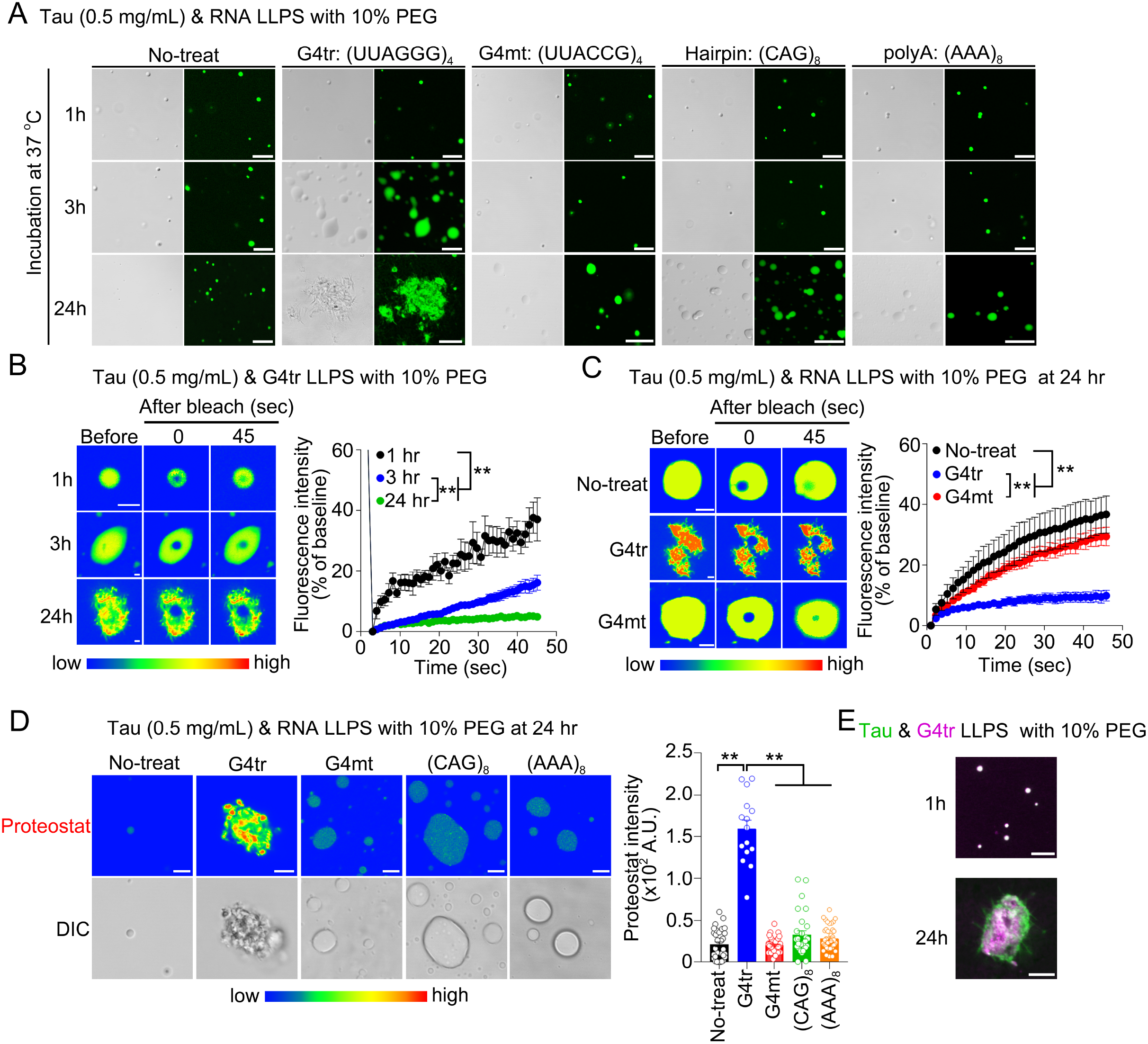
rG4 accelerates Tau liquid–solid phase transition. (A) Representative images of *in vitro* Tau (0.5 mg/mL) phase separation with 10% PEG and 1 μM RNA nucleotides at 1, 3, and 24 h after incubation. Scale bar, 5 μm. (B) FRAP assays of Tau LLPS with 10% PEG and 1 μM G4tr at 1, 3, and 24 h after incubation. Scale bar, 5 μm. n = 7 (1 h), n = 7 (3 h), and n = 6 (24 h). (C) FRAP assays of Tau LLPS with 10% PEG in the presence of 1 μM G4tr or G4mt at 24 h after incubation. Scale bar, 2 μm. n = 7 (1 h), n = 7 (3 h), and n = 6 (24 h). (D) Analysis of Proteostat intensity within Tau LLPS with 10% PEG and 1 μM RNA at 24 h after incubation. Scale bar, 5 μm. n = 31 (untreated), n = 15 (G4tr), n = 40 (G4mt), n = 28 ((CAG)_8_), and n = 40 ((AAA)_8_). (E) Representative images of Tau (green) and G4tr (magenta) with 10% PEG incubated at 37L for 1 and 24 h. Data are presented as the mean ± standard error of the mean. \*\**P* < 0.01 by two-way (B and C) and one-way (D) analysis of variance with Bonferroni’s multiple comparisons test.

To investigate whether Tau preferentially interacts with rG4 compared to other RNA structures, we performed surface plasmon resonance (SPR) binding analysis. The SPR analysis showed no significant differences in the affinity for any RNA structure, although the affinity of Tau for rG4 appeared to be slightly higher than that for other RNA structures (**Table. 1; Fig. S2**).

### Ca^2+^ facilitates Tau LLPS and rG4-induced Tau phase transition

Persistent and excessive Ca^2+^ entry in neurons could be a major factor in the development of neurodegeneration, including tauopathies [22, 23]. In addition, Tau aggregation is facilitated by divalent cations, including Ca^2+^ *in vitro* [24, 25]. Thus, we investigated whether Ca^2+^ affects Tau LLPS- and rG4-induced Tau phase transitions *in vitro*. We found that Ca^2+^ promoted Tau phase separation when incubated at 37 □ after 1 h (**Fig. 3A**). Moreover, fluorescence recovery of fluorescein-labeled Tau in FRAP was markedly reduced in the presence of Ca^2+^ (500 μM) after 24 h (**Fig. 3B**), suggesting Ca^2+^ may be a key cofactor of Tau phase transition.

**Fig. 3.**
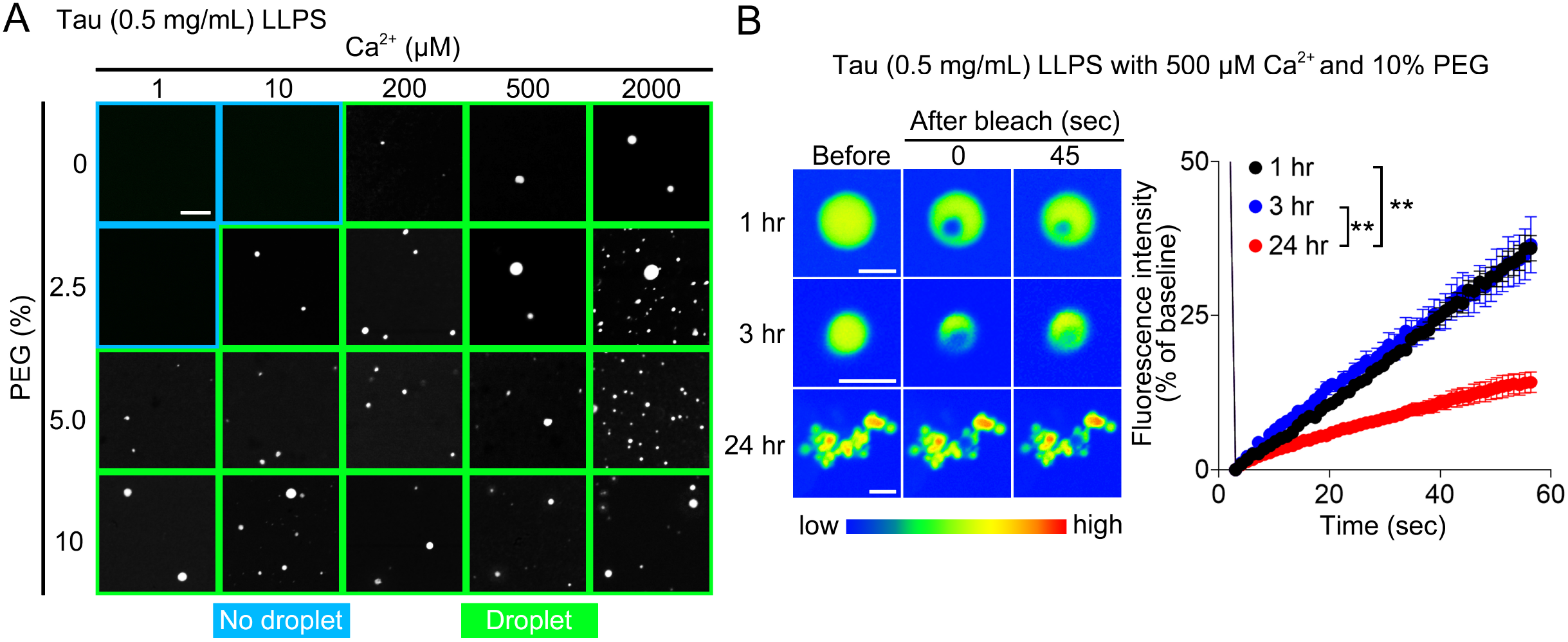
Ca^2+^ promotes Tau LLPS and liquid–solid phase transition. (A) Representative images of *in vitro* Tau (0.5 mg/mL) phase separation dependent on PEG and Ca^2+^ concentrations when incubated at 37L for 1 h. Scale bar, 5 μm. (B) FRAP assays of Tau LLPS with 10% PEG at 1, 3, and 24 h after incubation. Scale bar, 2 μm. n = 9 per time point. Data are presented as the mean ± standard error of the mean. \*\**P* < 0.01 by two-way (B) analysis of variance with Bonferroni’s multiple comparisons test.

Notably, Cyanine5-labeled G4tr co-aggregated with fluorescein-labeled Tau in the presence of Ca^2+^ and 10% PEG when incubated at 37L for 1 h (**Fig. 4A**). Ca^2+^ accelerated Tau phase transition by G4tr, resulting in a significant increase in proteostatic intensity and a decrease in Tau fluorescence recovery in FRAP, which was prevented by EGTA (**Fig. 4B and 4C**). Conversely, Ca^2+^ did not affect the G4mt-Tau complex with 10% PEG (**Fig. S3**). Finally, even under PEG-free conditions, Ca^2+^ induced the G4tr-Tau complex to shift to an aggregate-like state after 1 h, thereby significantly increasing the proteostat intensity, whereas this effect was attenuated by EGTA (**Fig. 4D and 4E**).

**Fig. 4.**
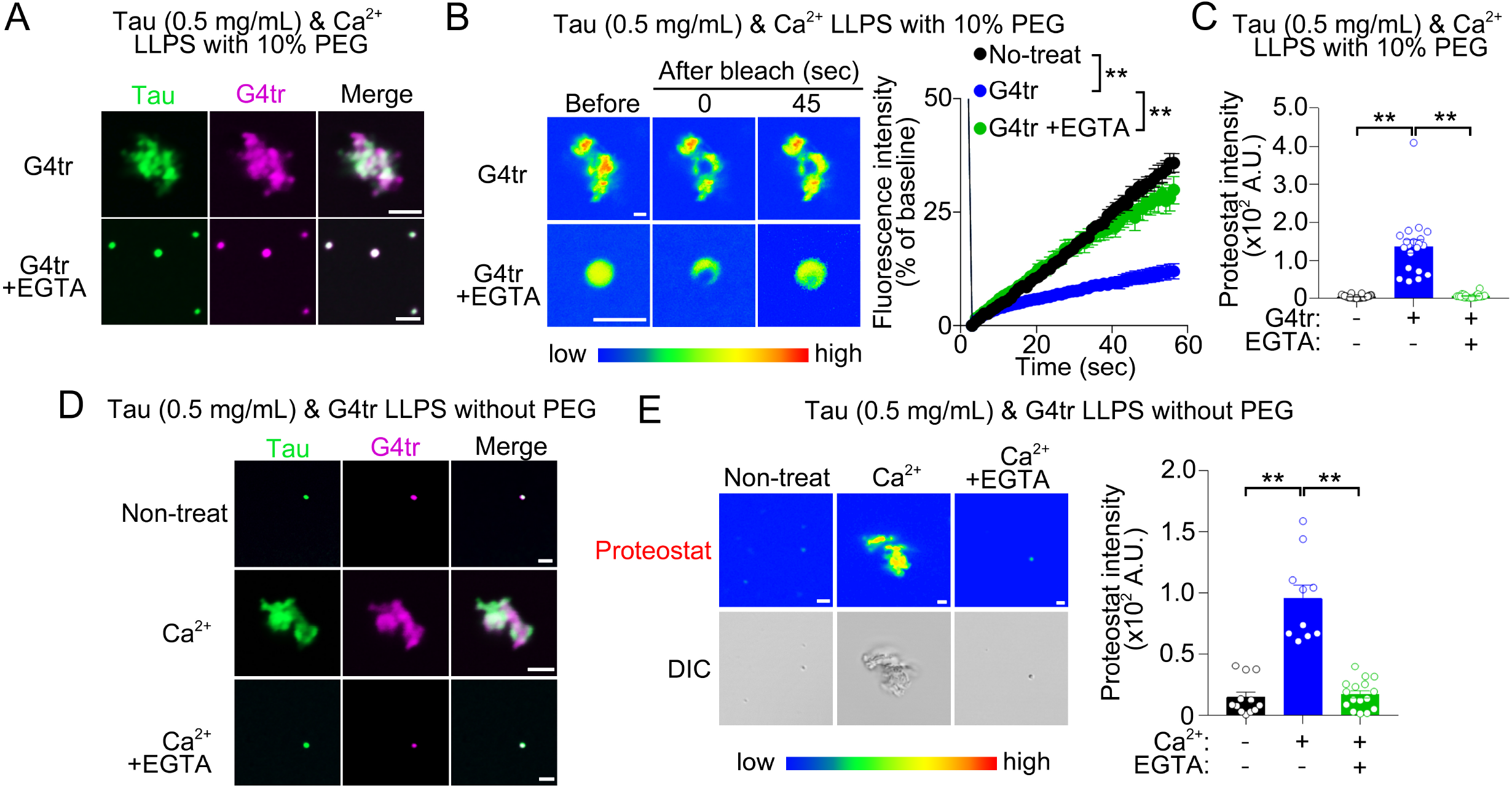
Synergistic effect of Ca^2+^ and G4tr on Tau liquid–solid phase transition. (A) Representative images of *in vitro* Tau (0.5 mg/mL; green) and G4tr (1 μM; magenta) phase separation in the presence of 10% PEG and 500 μM Ca^2+^ with or without 2.5 mM EGTA when incubated at 37L for 1 h. Scale bar, 2 μm. (B) FRAP assays of Tau LLPS in the presence of 10% PEG, 1 μM G4tr, and 500 μM Ca^2+^ with or without 2.5 mM EGTA when incubated at 37L for 1 h. Scale bar, 2 μm. n = 9 (untreated), n = 8 (G4tr), and n = 6 (G4tr + EGTA). (C) Analysis of Proteostat intensity within Tau LLPS with 10% PEG, 1 μM G4tr, and 500 μM Ca^2+^ with or without 2.5 mM EGTA when incubated at 37L for 1 h. Scale bar, 5 μm. n = 35 (untreated), n = 20 (G4tr), and n = 22 (G4tr + EGTA). (D) Representative images of *in vitro* Tau (0.5 mg/mL; green) and G4tr (1 μM; magenta) phase separation with or without 500 μM Ca^2+^ and 2.5 mM EGTA when incubated at 37L for 1 h. Scale bar, 2 μm. (E) Analysis of Proteostat intensity within Tau LLPS with 1 μM G4tr in the presence or absence of 500 μM Ca^2+^ and 2.5 mM EGTA when incubated at 37L for 1 h. Scale bar, 2 μm. n = 12 (G4tr), n = 10 (G4tr + Ca^2+^), and n = 16 (G4tr + Ca^2+^ + EGTA). Data are presented as the mean ± standard error of the mean. \*\**P* < 0.01 by two-way (B) and one-way (C and E) analysis of variance with Bonferroni’s multiple comparisons test.

## Discussion

In the present study, we demonstrated that rG4 promoted the liquid-to-solid phase transition of Tau *in vitro*. Cell-derived total RNAs transform Tau droplets from a liquid to a gel-like state. We found that rG4 in RNAs enables the phase transition of Tau. Moreover, Ca^2+^ can facilitate Tau phase separation and aggregation, thereby accelerating rG4-induced Tau phase transition. Taken together, these results suggest that rG4 and Ca^2+^ may be key cofactors in Tau aggregation.

Although our study showed heparin-enlarged Tau droplets without phase transition under physiological conditions (**Fig. 1**), previous reports have demonstrated that heparin induces Tau aggregation *in vitro* under non-physiological conditions [14, 17, 19, 26]. Elevated cationic strength (> 50 mM NaCl) significantly attenuates heparin-induced Tau aggregation [27-29], suggesting that decreased electrostatic interactions by mimicking intracellular ion conditions (140 mM KCl, 15 mM NaCl, and 10 mM MgCl_2_) may prevent the effect of heparin on Tau aggregation [27]. In addition, the aggregated form of Tau induced by heparin differs from that in tauopathies, including AD and frontotemporal dementia [30, 31]. Moreover, heparin-induced Tau aggregates fail to initiate Tau aggregation in primary neurons and mouse brains [32]. These data suggest that the aggregation of Tau by polyanions may be artificial and not occur in cells.

Although rG4 facilitated the Tau liquid-to-solid phase transition (**Fig. 2**), there were no dramatic changes in its affinity for Tau compared to that of other structural RNAs (**Table 1; Fig. S2**). The process of LLPS in prion-like RNA-binding proteins (RBPs), including Tau, is differentially influenced by the type of RNA [33-36], suggesting that the specific domain of RBPs to which the RNA binds and the characteristics of the RNA, such as length and structure, rather than the binding affinity of the RNA, intricately affect Tau LLPS [35]. For example, fused in sarcoma (FUS), a representative amyotrophic lateral sclerosis (ALS)-linked RBP, undergoes a liquid-to-solid phase transition by adding rG4, but not randomized RNAs [37]. Other ALS-linked RBP and TAR DNA-binding protein 43 (TDP-43) condensates are buffered by the addition of tRNA, whereas nuclear-enriched abundant transcript 1 lncRNA, which can form rG4, facilitates TDP-43 LLPS [34, 38]. Moreover, rG4 within prion mRNA may contribute to destabilizing the structure of prion proteins and trigger their conversion to an infectious form [39]. We have also demonstrated that rG4 may change the structure of αSyn by binding to its N-terminus, leading to αSyn aggregation and in turn neurodegeneration in mouse neurons [21]. The effect of rG4 on the structure of prion-like RBPs and the mechanism by which rG4 initiates aggregation are yet to be resolved, suggesting that rG4 can alter these properties.

**Table 1.** Affinity of Tau for 24-mer RNA oligonucleotides as revealed by OpenSPR.

| Affinity of Tau-RNA |  |  |  |
| --- | --- | --- | --- |
| RNA | $K_a$ ($\times 10^2$ 1/s) | $K_d$ ($\times 10^{-4}$ 1/s) | $K_D$ (nM) |
| G4tr | $275 \pm 2.0$ | $5.39 \pm 0.52$ | $19.6 \pm 2.0$ |
| G4mt | $162 \pm 0.65$ | $5.11 \pm 0.18$ | $31.5 \pm 1.2$ |
| (CAG)<sub>8</sub> | $183 \pm 0.84$ | $6.53 \pm 0.025$ | $35.7 \pm 0.3$ |
| (AAA)<sub>8</sub> | $185 \pm 0.22$ | $6.5 \pm 0.039$ | $35.1 \pm 0.25$ |

In addition, we found that Ca^2+^ accelerated rG4-induced Tau phase transition (**Fig. 3 and 4**). Consistent with our observations, previous reports have indicated that Ca^2+^ can accelerate the fibril formation of Tau *in vitro* [25, 40]. Another bivalent cation, Zn^2+^, also facilitates Tau LLPS and fibrilization by binding to Cys residues present in the microtubule-binding repeat domain (Cys-291 and Cys-322) [25, 41], suggesting that Ca^2+^ binding to Cys in the microtubule-binding repeat domain may be associated with Tau aggregation [25]. Further studies are required to uncover the mechanism underlying rG4/Ca^2+^-induced Tau aggregation, such as the identification of the binding sites of rG4 and Ca^2+^ to Tau using nuclear magnetic resonance spectrometry.

Impaired RNA metabolism has been recognized as a common event in neurodegeneration, especially the formation of RNA granules, termed stress granules (SGs), which appear to be particularly relevant to neurodegenerative diseases, including tauopathy [42]. Cellular stress, such as treatment with MG-132 or arsenite, triggers the formation of TIA1-positive SGs; in turn, Tau is recruited to SGs in Tau-overexpressing mouse neuronal cell lines and primary neurons, resulting in an increase in misfolded and phosphorylated Tau [43, 44]. TIA1 knockdown attenuates Tau aggregation, neuronal death, and memory deficits in mouse models expressing human mutated Tau (P301L or P301S) [44, 45], suggesting that the recruitment of Tau into SGs may trigger aggregation. Kharel et al. reported that various cellular stressors markedly elevated rG4 levels in human cells [46]. Moreover, we previously demonstrated that rG4s are highly enriched in SGs and organize TIA1-positive SGs assemblies in primary mouse neurons [20]. These observations suggest that co-aggregation of rG4s and Tau in SGs under cellular stress conditions contributes to neurodegeneration. There has also been an increasing number of studies focusing on the role of rG4 in human aging and neurodegenerative diseases [47, 48]. In AD neurons, the accumulation of rG4 structures is observed in lamina-associated domains [49]. Importantly, Kallweit et al. demonstrated a significant positive correlation between rG4 accumulation and the Braak stage in the outer molecular layer region of the AD brain [50]. They also indicated that NFTs contain rG4s in AD neurons [50], suggesting that rG4s may contribute to Tau aggregation in the human brain.

In conclusion, rG4 and Ca^2+^ synergistically induced Tau aggregation *in vitro*. The application of heparin and other structured RNAs failed to transition Tau coacervates to an aggregate-like state, indicating that the rG4-induced Tau phase transition does not occur through electrostatic interactions. Since rG4 can initiate the phase transition of prion-like proteins including αSyn and FMRpolyG [20, 21], rG4 may be a common key factor in neurodegeneration.

### Experimental procedures

#### Purification and fluorescein-labeling of Tau proteins

The pET23a-human Tau441 plasmid was expressed in *Escherichia coli* BLR (DE3) cells harboring an overproducing plasmid and purified as previously described [26]. Briefly, the bacterial lysate obtained by ultrasonication was incubated at 80°C for 10 min, immediately cooled on ice, and centrifuged to remove insoluble matter. Streptomycin sulfate (final concentration, 2.5%) was added to the supernatant to precipitate the nucleic acids. After centrifugation, the supernatant was dialyzed overnight against a purification buffer (50 mM Tris-HCl buffer, 2 mM EDTA, pH 7.8). The dialysate was fractionated using cation-exchange chromatography (HiTrap SP HP; GE Healthcare). All chromatography steps were performed on an AKTA-explorer system (GE Healthcare) at 4°C. Samples were desalted and stored in a lyophilized state at 4°C until being assayed. For fluorescence detection, recombinant Tau proteins were labeled with fluorescein using the Fluorescein Labeling Kit-NH2, according to the manufacturer’s instructions (Dojindo Molecular Technologies Inc.).

#### RNA oligonucleotides

The RNA oligonucleotides were synthesized by Hokkaido System Science: non-labeled and biotin-labeled: r(UUAGGG)_4_, r(UUACCG)_4_, r(CAG)_8_, and r(AAA)_8_; Cyanine5-labeled: r(UUAGGG)_4_ and r(UUACCG)_4_. Each RNA was resolved in RNase-free water at a concentration of 100 μM.

#### *In vitro* LLPS, Proteostat, and FRAP assay

Non-labeled and/or fluorescein-labeled human Tau (0.005–0.5 mg/mL; 0.109–10.9 μM) were prepared in the LLPS assay buffer: 50 mM Tris-HCl buffer (pH 7.5) containing 140 mM KCl, 15 mM NaCl, 10 mM MgCl_2_, 0–10% PEG8000 (MP Biomedicals), and 1 unit/μL RNase inhibitor (TOYOBO), in the presence or absence of 0.001–2 mM CaCl_2_ and 2.5 mM EGTA. Heparin (0–10%) (NACALAI TESQUE, INC.), total RNA (1–100 ng/μL), or synthesized 24-mer RNA oligonucleotides (1 μM) were added and incubated at 37× for 1, 3, and/or 24 h. Total RNA was extracted using an RNeasy Mini Kit (Qiagen) from intact Neuro-2a cells. Some samples with total RNA (in the absence of an RNase inhibitor) were treated with RNase A (8 μg/μL; Qiagen) or Cryonase Cold-active Nuclease (1.6 U/μL; TAKARA) for 1 h before adding the Tau protein. Samples were incubated at 37× for 1, 3, or 24 h.

Tau LLPS was also investigated under non-physiological conditions (25 mM HEPES buffer pH 7.4 in 10 mM NaCl and 10% PEG at room temperature (25_±_2 □); 30 mM Tris-HCl buffer pH 7.5 in 10 mM NaCl, 15% PEG and 3% heparin with 200 rpm agitation at 37 □; 20 mM HEPES buffer pH 7.0 in 17% heparin at room temperature (25 _±_2 □)) at 1 and 24 h after incubation according to previous reports [11, 14, 19]. The reacted solutions were mounted on glass slides using a 0.12 nm spacer (Sigma-Aldrich) and a coverslip. For fluorescence observations, non-labeled human Tau proteins were mixed with fluorescein-labeled proteins at a ratio of 1:1. To assess the aggregated form of Tau, 2% (v/v) proteostat reagent (Enzo Life Sciences) was added to the LLPS assay buffer. Differential interference contrast images were obtained using a TCS SP8 confocal microscope (Leica Microsystems). Protein intensities were analyzed using the LAS X system (Leica Microsystems). For the FRAP assay, samples were photobleached with 50% laser power, and time-lapse images were recorded every 1 s using a Zeiss Objective Plan-Apochromat 63×/1.4 oil DIC M27 to track photorecovery behavior using an LSM900 microscopy system (Carl Zeiss).

#### SPR-binding assays

SPR experiments were performed using OpenSPR (Nicoya) as previously reported [21]. Briefly, biotin-labeled RNA oligonucleotides were fixed with a sensor chip using a biotin-streptavidin sensor kit (Nicoya), and Tau in running buffer (50 mM Tris-HCl pH 7.5 and 10 mM KCl) was streamed over the sensor chip to allow interaction. After ligand signal stabilization, the running buffer flowed at a rate of 20 μL/min for 5 min to collect dissociation data. Binding kinetic parameters were obtained by fitting the curve to a one-to-one binding model using TraceDrawer software (Ridgeview Instruments).

## Statistical analysis

Data are presented as the mean ± standard error of the mean. The statistical significance of differences among groups was tested by one-way (proteostat intensity) or two-way analysis (FRAP assay) of variance with *post-hoc* Bonferroni’s multiple comparison test. *P* < 0.05 represented a statistically significant difference. All statistical analyses were performed using the GraphPad Prism 7 software (GraphPad Software).

## Data availability

All data needed to evaluate the conclusions in the paper are present in the paper. Additional data related to this paper may be requested from the corresponding author upon reasonable request.

Corresponding author: Yasushi Yabuki and Norifumi Shioda

(to Y.Y.)

(to N.S.)

## Acknowledgments

We thank editage for the English language review.

## Author Contributions

Y.Y. and N.S. designed the study. Y.Y., K.M., G.K., K.K., K.H., and S.I. performed the experiments. Y. K. and T. M. provided critical advice on the synthesis and purification of Tau. Y.Y. and N.S. wrote the manuscript. Y.Y. and N.S. supervised the study.

## Corresponding author

Correspondence should be addressed to Yasushi Yabuki and Norifumi Shioda.

## Funding and additional information

This work was supported by AMED (grant numbers: JP21wm0525023 and JP21gm6410021 to Y.Y.), JSPS KAKENHI (grant numbers: JP21K06579 and JP23H03851 to Y.Y.; JP21K20723 and JP22J00687 to K.M.; and 21H00207, 20K21400, and JP20H03393 to N.S.), JST FOREST Program (JPMJFR2043 to N.S.), Narishige Neuroscience Research Foundation (to Y.Y.), International Research Core for Stem Cell-based Developmental Medicine Encourage Project, Institute of Molecular Embryology and Genetics, Kumamoto University.

## Reference

1. Lee, G., Cowan, N., and Kirschner, M. (1988) The primary structure and heterogeneity of tau protein from mouse brain. Science. 239(4837), 285–288

2. Weingarten, M. D., Lockwood, A. H., Hwo, S. Y., and Kirschner, M. W. (1975) A protein factor essential for microtubule assembly. Proc. Natl. Acad. Sci. 72, 1858–1862

3. Vacchi, E., Kaelin-Lang, A., and Melli, G. (2020) Tau and alpha synucleins synergistically effect neurodegenerative diseases when the periphery is the core. Int. J. Mol. Sci. 21(14), 5030

4. Barghorn, S., Zheng-Fischhofer, Q., Ackmann, M., Biernat, J., von Bergen, M., Mandelkow, E. M., et al. (2000) Structure, microtubule interactions, and paired helical filament aggregation by tau mutants of frontotemporal dementias. Biochem. 39, 11714–11721

5. Hong, M., Zhukareva, V., Vogelsberg-Ragaglia, V., Wszolek, Z., Reed, L., Miller, B. I., et al. (1998) Mutation-specific functional impairments in distinct tau isoforms of hereditary FTDP-17. Science. 282, 1914–1917

6. Guan, P. P., and Wang, P. (2023) The involvement of post-translational modifications in regulating the development and progression of Alzheimer’s Disease. Mol. Neurobiol. 60(7), 3617–3632

7. Brangwynne, C. P., Eckmann, C. R., Courson, D. S., Rybarska, A., Hoege, C., Gharakhani, J., et al. (2009) Germline P granules are liquid droplets that localize by controlled dissolution/condensation. Science. 324, 1729–1732

8. Jain, A., and Vale, R. D. (2017) RNA phase transitions in repeat expansion disorders. Nature. 546, 243–247

9. Alberti S., Gladfelter, A., and Mittag, T. (2019) Considerations and challenges in studying liquid–liquid phase separation and biomolecular condensates. Cell. 176, 419–434

10. Hernández-Vega, A., Braun, M., Scharrel, L., Jahnel, M., Wegmann, S., Hyman, B. T., et al. (2017) Local nucleation of microtubule bundles based on tubulin concentration in the condensed tau phase. Cell Rep. 20(10), 2304–2312

11. Hochmair, J., Exner, C., Franck, M., Dominguez-Baquero, A., Diez, L., Brognaro, H., et al. (2022) Molecular crowding and RNA synergize to promote phase separation, microtubule interaction, and seeding of Tau condensates. EMBO J. 41(11), e108882

12. Wegmann, S., Eftekharzadeh, B., Tepper, K., Zoltowska, K. M., Bennett, R. E., Dujardin, S., et al. (2018) Tau protein liquid-liquid phase separation can initiate tau aggregation. EMBO J. 37(7), e98049

13. Kanaan, N. M., Hamel, C., Grabinski, T., and Combs, B. (2020) Liquid-liquid phase separation induces pathogenic tau conformations in vitro. Nat. Commun. 11(1), 2809

14. Wen, J., Hong, L., Krainer, G., Yao, Q. Q., Knowles, T. P. J., Wu, S., et al. (2021) Conformational expansion of tau in condensates promotes irreversible aggregation. J. Am. Chem. Soc. 143(33), 13056–13064

15. Boyko, S., and Surewicz, W. K. (2022) Tau liquid-liquid phase separation in neurodegenerative diseases. Trends Cell Biol. 32(7), 611–623

16. Ukmar-Godec, T., Hutten, S., Grieshop, M. P., Rezaei-Ghaleh, N., Cima-Omori, M.S., Biernat, J., et al. (2019) Lysine/RNA-interactions drive and regulate biomolecular condensation. Nat. Commun. 10(1), 2909

17. Boyko, S., Surewicz, K., and Surewicz, W. K. (2020) Regulatory mechanisms of tau protein fibrillation under the conditions of liquid-liquid phase separation. Proc. Natl. Acad. Sci. 117(50), 31882–31890

18. Sen, D., and Gilbert, W. (1988) Formation of parallel 4-stranded complexes by guanine-rich motifs in DNA and its implications for meiosis. Nature. 334, 364–366

19. Lin, Y., Fichou, Y., Zeng, Z., Hu, N. Y., and Han, S. (2020) Electrostatically driven complex coacervation and amyloid aggregation of Tau are independent processes with overlapping conditions. ACS Chem. Neurosci. 11(4), 615–627

20. Asamitsu, S., Yabuki, Y., Ikenoshita, S., Kawakubo, K., Kawasaki, M., Usuki, S., et al. (2021) CGG repeat RNA G-quadruplexes interact with FMR polyG to cause neuronal dysfunction in fragile X-related tremor/ataxia syndrome. Sci. Adv. 7, eabd9440

21. Matsuo, K., Asamitsu, S., Maeda, K., Kawakubo, K., Komiya, G., Kudo, K., et al. (2023) RNA G-quadruplexes forming scaffolds for α-synuclein aggregation lead to progressive neurodegeneration. bioRxiv. 548322

22. Webber, E. K., Fivaz, M., Stutzmann, G. E., and Griffioen, G. (2023) Cytosolic calcium: judge, jury, and executioner of neurodegeneration in Alzheimer’s disease and beyond. Alzheimers Dement. 19(8), 3701–3717

23. Imamura, K., Sahara, N., Kanaan, N. M., Tsukita, K., Kondo, T., Kutoku, Y., et al. (2016) Calcium dysregulation contributes to neurodegeneration in FTLD patient iPSC-derived neurons. Sci. Rep. 6, 34904

24. Singh, V., Xu, L., Boyko, S., Surewicz, K., and Surewicz, W. K. (2020) Zinc promotes liquid-liquid phase separation of the tau protein. J. Biol. Chem. 295(18), 5850–5856

25. Moreira, G.G., Cristóvão, J.S., Torres, V.M., Carapeto, A.P., Rodrigues, M.S., Landrieu, I., et al. (2019). Zinc Binding to Tau Influences Aggregation Kinetics and Oligomer Distribution. Int J Mol Sci. 20(23), 5979.

26. Ogawa, K., Ishii, A., Shindo, A., Hongo, K., Mizobata, T., Sogon, T., et al. (2020) Spearmint extract containing rosmarinic acid suppresses amyloid fibril formation of proteins associated with dementia. Nutrients. 12(11), 3480

27. Friedhoff, P., Schneider, A., Mandelkow, E. M., and Mandelkow, E. (1998) Rapid assembly of Alzheimer-like paired helical filaments from microtubule-associated protein tau monitored by fluorescence in solution. Biochem. 37(28), 10223–10230

28. Jeganathan, S., von Bergen, M., Mandelkow, E. M., and Mandelkow, E. (2008) The natively unfolded character of tau and its aggregation to alzheimer-like paired helical filaments. Biochem. 47(40), 10526–10539

29. Lin, Y., Fichou, Y., Longhini, A. P., Llanes, L. C., Yin, P., Bazan, G. C., et al. (2021) Liquid-liquid phase separation of Tau driven by hydrophobic interaction facilitates fibrillization of Tau. J. Mol. Biol. 433(2), 166731

30. Zhang, W., Falcon, B., Murzin, A. G., Fan, J., Crowther, R. A., Goedert M., et al. (2019) Heparin-induced tau filaments are polymorphic and differ from those in Alzheimer’s and Pick’s diseases. Elife. 8, e43584

31. Zeng, Y., Yang, J., Zhang, B., Gao, M., Su, Z., and Huang, Y. (2021) The structure and phase of tau: from monomer to amyloid filament. Cell. Mol. Life Sci. 78(5), 1873–1886

32. Guo, J. L., Narasimhan, S., Changolkar, L., He, Z., Stieber, A., Zhang, B., et al. (2016) Unique pathological tau conformers from Alzheimer’s brains transmit tau pathology in nontransgenic mice. J. Exp. Med. 213(12), 2635–2654

33. Zhang, X., Lin, Y., Eschmann, N. A., Zhou, H., Rauch, J. N., Hernandez, I., et al. (2017) RNA stores tau reversibly in complex coacervates. PLoS Biol. 15(7), e2002183

34. Wang, C., Duan, Y., Duan, G., Wang, Q., Zhang, K., Deng, X., et al. (2020) Stress induces dynamic, cytotoxic antagonizing TDP-43 nuclear bodies via paraspeckle lncRNA NEAT1-mediated liquid-liquid phase separation. Mol. Cell. 79(3), 443–458

35. Portz, B., Lee, B. L., and Shorter, J. (2021) FUS and TDP-43 phases in health and disease. Trends Biochem. Sci. 46(7), 550–563

36. Hurtle, B. T., Xie, L., and Donnelly, C. J. (2023) Disruption of pathological phase transitions during neurodegeneration. J. Clin. Invest. 133(13), e168549

37. Ishiguro, A., Lu, J., Ozawa, D., Nagai, Y., and Ishihama, A. (2021) ALS-linked FUS mutations dysregulate G-quadruplex-dependent liquid-liquid phase separation and liquid-to-solid transition. J. Biol. Chem. 297(5), 101284

38. Simko, E. A. J., Liu, H., Zhang, T., Velasquez, A., Teli, S., Haeusler, A. R., et al. (2020) G-quadruplexes offer a conserved structural motif for NONO recruitment to NEAT1 architectural lncRNA. Nucleic Acids Res. 48(13), 7421–7438

39. Olsthoorn, R. C. (2014) G-quadruplexes within prion mRNA: the missing link in prion disease? Nucleic Acids Res. 42(14), 9327–9233

40. Yang, L. S., and Ksiezak-Reding, H. (1999) Ca2+ and Mg2+ selectively induce aggregates of PHF-tau but not normal human tau. J. Neurosci. Res. 55(1), 36–43.

41. Mo, Z. Y., Zhu, Y. Z., Zhu, H. L., Fan, J. B., Chen, J., and Liang, Y. (2009) Low micromolar zinc accelerates the fibrillization of human tau via bridging of Cys-291 and Cys-322. J. Biol. Chem. 284(50), 34648–34657

42. Wolozin, B., and Ivanov, P. (2019) Stress granules and neurodegeneration. Nat. Rev. Neurosci. 20(11), 649–666

43. Piatnitskaia, S., Takahashi, M., Kitaura, H., Katsuragi, Y., Kakihana, T., Zhang, L., et al. (2019) USP10 is a critical factor for Tau-positive stress granule formation in neuronal cells. Sci. Rep. 9(1), 10591

44. Vanderweyde, T., Apicco, D. J., Youmans-Kidder, K., Ash, P. E. A., Cook, C., Lummertz da Rocha, E., et al. (2016) Interaction of tau with the RNA-binding protein TIA1 regulates tau pathophysiology and toxicity. Cell Rep. 15(7), 1455–1466

45. Apicco, D. J., Ash, P. E. A., Maziuk, B., LeBlang, C., Medalla, M., Al Abdullatif, A., et al. (2018) Reducing the RNA binding protein TIA1 protects against tau-mediated neurodegeneration in vivo. Nat. Neurosci. 21(1), 72–80

46. Kharel, P., Fay, M., Manasova, E. V., Anderson, P. J., Kurkin, A. V., Guo, J. U., et al. (2023) Stress promotes RNA G-quadruplex folding in human cells. Nat. Commun. 14(1), 205.

47. Vijay Kumar, M. J., Morales, R., and Tsvetkov, A. S. (2023) G-quadruplexes and associated proteins in aging and Alzheimer’s disease. Front. Aging. 4, 1164057

48. Antcliff, A., McCullough, L. D., and Tsvetkov, A. S. (2021) G-Quadruplexes and the DNA/RNA helicase DHX36 in health, disease, and aging. Aging. 13(23), 25578–25587

49. Hanna, R., Flamier, A., Barabino, A., and Bernier, G. (2021) G-quadruplexes originating from evolutionary conserved L1 elements interfere with neuronal gene expression in Alzheimer’s disease. Nat. Commun. 12(1), 1828

50. Kallweit, L., Hamlett, E. D., Saternos, H., Gilmore, A., Granholm, A. C., and Horowitz, S. (2023) A new role for RNA G-quadruplexes in aging and alzheimer’s disease. bioRxiv. 10.02.560545581078v1

