## supplementary information for "RNA G-quadruplexes and calcium ions synergistically induce Tau phase transition *in vitro*"

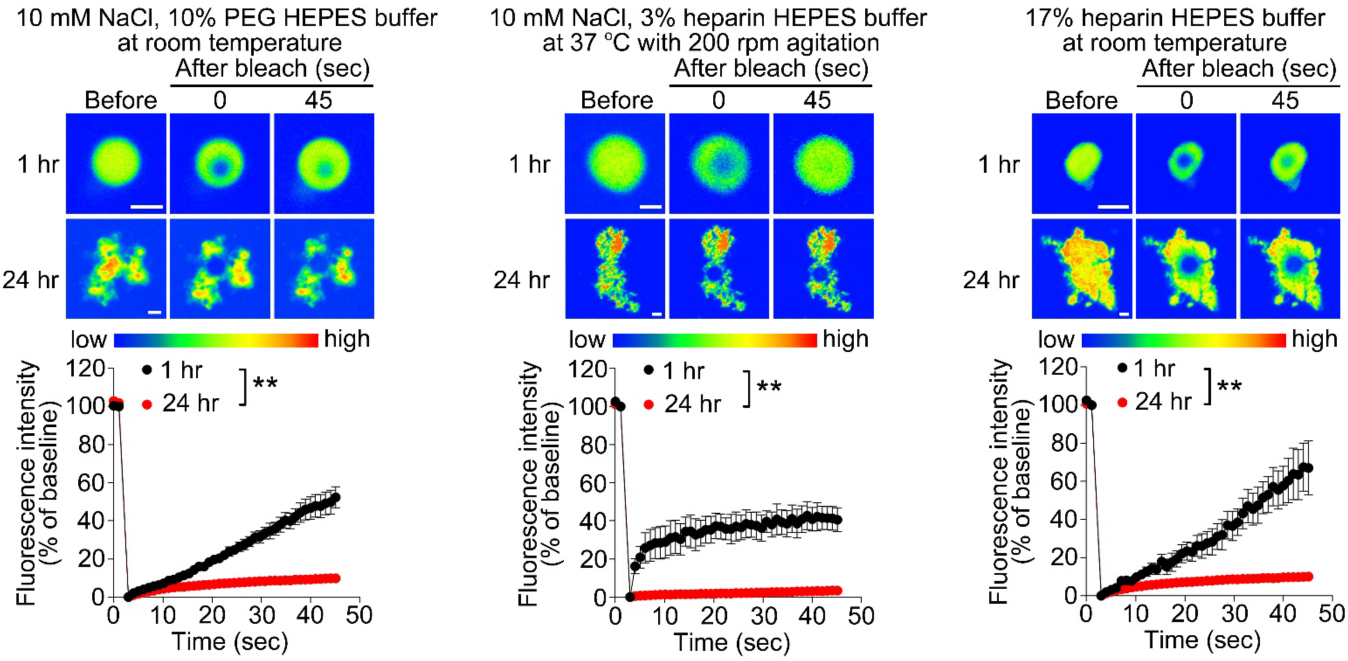


**Fig. S1 Tau LLPS at non-physiological conditions.**

FRAP assays of Tau LLPS incubated for 1 or 24 h at non-physiological conditions: 25 mM HEPES buffer pH 7.4 with 10 mM NaCl and 10% PEG at room temperature (25 ± 2°C) (right); 30 mM Tris-HCl buffer pH 7.5 with 10 mM NaCl, 15% PEG, and 3% heparin with 200 rpm agitation at 37°C (center); 20 mM HEPES buffer pH 7.0 with 17% heparin at room temperature (25 ± 2°C) (left). n = 5–7 per group per time point. Data are presented as the mean ± standard error of the mean. **$P < 0.01$ by two-way analysis of variance with Bonferroni's multiple comparisons test.

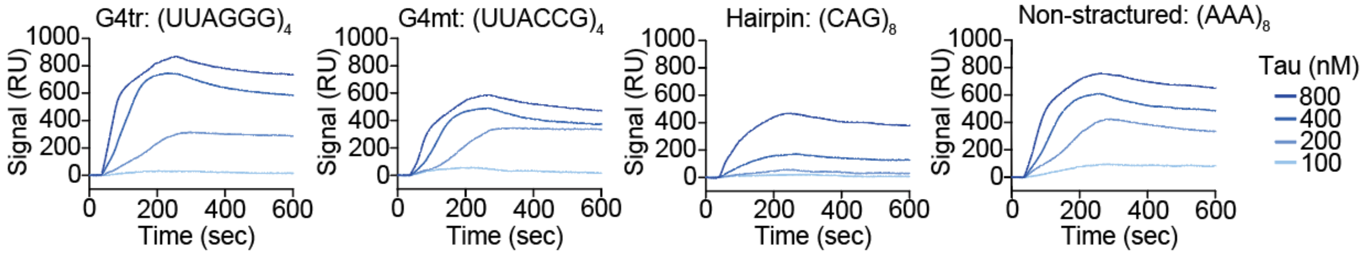


**Fig. S2 SPR sensorgrams for the interaction of Tau with 24-mer RNA oligonucleotides.**

The SPR sensorgrams used in the analysis for Table 1.

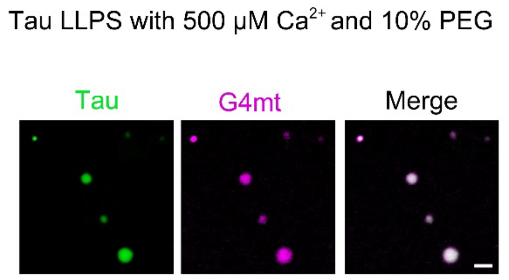


**Fig. S3 $Ca^{2+}$ does not affect Tau and G4mt condensates.**

Representative images of *in vitro* Tau (0.5 mg/mL; green) and G4mt (1 µM; magenta) phase separation in the presence of 10% PEG and 500 µM $Ca^{2+}$ when incubated at 37°C for 1 h. Scale bar, 2 µm.
